# Differential occupancy and regulatory interactions of KDM6A in bladder cell lines

**DOI:** 10.1101/2022.12.19.520968

**Authors:** Gülden Özden-Yılmaz, Büşra Savaş, Ahmet Bursalı, Aleyna Eray, Alirıza Arıbaş, Şerif Şentürk, Ezgi Karaca, Gökhan Karakülah, Serap Erkek-Ozhan

## Abstract

Epigenetic deregulation is a critical theme which needs further investigation for bladder cancer research. One of the highly mutated genes in bladder cancer is *KDM6A*, functioning as a H3K27 demethylase and is part of the MLL3/4 complexes. To decipher the role of KDM6A in normal versus tumor setting, we identified the genomic landscape of KDM6A in normal, immortalized and cancer bladder cells. Our results showed differential KDM6A occupancy at the genes involved in cell differentiation, chromatin organization and Notch signaling depending on the cell type and the mutation status of *KDM6A*. Transcription factor motif analysis revealed HES1 to be enriched at KDM6A peaks identified for T24 bladder cancer cell line, which has a truncating mutation in KDM6A, lacking demethylase domain. Our co-immunoprecipitation experiments reveal TLE co-repressors and HES1 as potential truncated and wild type KDM6A interactors. With the aid of structural modeling, we explored how the truncated KDM6A could interact with TLE, HES1, as well RUNX, HHEX transcription factors. These structures provide a solid mean to study the functions of KDM6A independent of its demethylase activity. Collectively, our work provides important contributions to the understanding of KDM6A malfunction in bladder cancer.

## 1. Introduction

Chromatin level-deregulation is regarded as one of the hallmarks of cancer, reflected at multiple different levels and affecting many processes driving tumorigenesis [1]. Malfunction in epigenetic regulatory mechanisms is especially noteworthy in bladder cancer considering the prominently high rate of mutations in chromatin modifier genes [2], which underscores the importance of studying epigenetic regulation in this cancer type.

Bladder cancer is one of the most common and lethal diseases worldwide. Each year, 430,000 patients are diagnosed with bladder cancer, 170,000 of which results in death [3]. 75-80% of bladder cancers show a non-muscle invasive character, while the remaining is muscle invasive [2]. Despite the recent genomic and molecular characterizations, cisplatin-based chemotherapy and radiotherapy-based approaches still remain the main treatment options for this cancer type [4]. Thus, within the context of high chromatin modifier mutations in this cancer [2], identification of epigenetic signatures implicated in pathogenesis of this disease is crucial.

Among the chromatin modifier mutations observed for bladder cancer, a few of them are especially high, including KMT2D (28%), KDM6A (26%), and ARID1A (25%), which are observed for muscle-invasive bladder cancer [2]. Mutations identified for KDM6A in bladder cancer have loss of function character [5] and there are many mutations identified within intrinsically disordered region (IDR) of KDM6A with one major mutation hotspot at position Q555 [5]. In this study, our focus is to identify the differential function of KDM6A in normal, immortalized, and cancer bladder cell lines. KDM6A is a H3K27 (lysine27 on histone H3) demethylase [6, 7], functioning as a crucial chromatin modifier required for various developmental processes and for cellular differentiation [8]. Importantly, KDM6A is mutated in the Kabuki syndrome, leading to facial abnormalities and cognitive dysfunction [8]. KDM6A is also required for embryonic development [9] and is involved in the resolution of bivalent chromatin domains [10]. KDM6A’s role in the regulation of *Hox* genes is well characterized [6]. In addition, KDM6A function has been identified to be important for the heart development [11] and hematopoiesis [12]. On the biochemical level, KDM6A has been shown to be associated with the evolutionarily conserved MLL3/MLL4 COMPASS complex, involved in the methylation of H3K4 and active chromatin organization in several cell lines [7, 8]. KDM6A is directly bound to the C-terminal of MLL3/MLL4 [13]. This indicates that KDM6A may play a role in regulating the catalytic activity of MLL3/4.

KDM6A is mutated in 26% of muscle-invasive bladder cancer cases, whereas its mutation frequency is higher in non-muscle invasive bladder cancer (47.6%) (cBioportal TCGA2017, MSK2017). However, the exact molecular mechanisms of how KDM6A functions in normal cellular setting and how it promotes tumorigenesis are currently open questions that need to be addressed in detail. Expanding on this, in this study, we generated KDM6A ChIP-seq data for one normal urothelial cell line (BdEC), one immortalized non-tumorigenic urothelial cell line (SV-HUC-1), and one urothelial cancer cell line (T24). The hypotheses we drove from the ChIP-seq data were further explored by biochemical assays and structural modeling. As a result, we show that KDM6A has a differential occupancy profile in the analyzed cell lines and it shows distinct behavior when it is mutated. We also identify the involvement of KDM6A in wide-spread regulation of Notch signaling in bladder cell lines. Furthermore, we structurally model the interaction of truncated KDM6A with repressors and co-repressors. Collectively, these results provide clear insights on previously uncovered roles of KDM6A in gene regulation.

## 2. Materials and Methods Experimental procedures

### 2.1. Cell culture

BdEC (primary bladder epithelial cells) (ATCC^®^ PCS-420-010^™^), SV-HUC-1 (Immortalized uroepithelial cells) (ATCC^®^ CRL-9520^™^), T24 (Urinary bladder transitional cell carcinoma) (ATCC^®^ HTB-4^™^) and HEK293T cell lines were used for experiments. BdEC cell line was grown in the appropriate medium (ATCC^®^ PCS-440-030^™^) and supplemented with the Prostate Epithelial Cell Growth Kit (ATCC PCS-440-040) as recommended by the ATCC and primocin (Invivogen). SV-HUC-1 was grown with RPMI-1640, T24 and HEK293T were grown with DMEM, supplemented with 10% FBS and 1% penicillin. Cells were maintained in a humidified incubator at 37°C and 5% CO_2_.

### 2.2. Chromatin Immunoprecipitation

Chromatin Immunoprecipitation assays were performed by following the protocol from Weber et al, 2007 [14]. Briefly, the cells were fixed with 1% formaldehyde for 15 minutes when they reached 70-80% confluence. Final concentration 0.125 M glycine was used to stop the crosslinking. After the pellet was lysed with Lysis Buffer, chromatin was fragmented using S220 Covaris Ultrasound Sonicator. With the intent of determining the optimum sonication for 200-500bp fragmented DNA, a 50 μl input sample was taken from the lysate. The rest of the chromatin was used for the immunoprecipitation (IP). For IP, chromatin was first pre-cleared with Dynabeads (Invitrogen, 11203D) blocked with BSA and tRNA and then bound with 5 μl KDM6A antibody (Cell Signaling, 33510) overnight at 4°C. Next day, chromatin-antibody complex were bound with pre-blocked Dynabeads for 3 hours at 4°C after the washing steps, chromatin was eluted. For both input and IP DNA, following the RNase and Proteinase K treatment, samples were incubated overnight at 65°C for reversal of crosslink. DNA was eluted using Zymo Research DNA Clean&Concentrator kit (D4034).

### 2.3. ChIP-seq

For T24, BdEC and SV-HUC-1 cell lines, ChIP and input DNA's were sent to EMBL, GeneCore facility for library preparation and sequencing. NEBNext DNA Ultra II kit was used for library preparation. Libraries were sequenced on Nextseq 500 platform using 75bp SE high output mode.

### 2.4. RT-qPCR

Total RNAs were isolated with MN Nucleospin RNA kit from each cell line. 0.25 μg RNA was used for cDNA conversion with Maxima First Strand cDNA Synthesis Kit. PCR was performed on the Applied Biosystems 7500 Fast real-time machine with Roche FastStart Essential DNA Green Master (SYBR) kit for the selected genes. Experiments were performed as three technical replicates.

### 2.5. ChIP-qPCR

After performing the KDM6A ChIP experiments for T24, SV-HUC-1, and BdEC cell lines, qPCR was performed using Roche FastStart Essential DNA Green Master (SYBR) kit on Applied Biosystems 7500 Fast real-time machine and the results were presented as % input IP. Experiments were performed as three technical replicates. ChIP-qPCR results were calculated using ΔΔct method. ChIP Ct values were normalized to the input.

### 2.6. Primer design

Primers for RT-qPCR and ChIP-qPCR were designed using primer designing tool Primer-Blast (NCBI). List of all the primers can be found in Table S3.

### 2.7. Western Blotting

Cells were washed with ice-cold PBS and scraped with 1x RIPA Buffer supplemented with protease inhibitor cocktail. After vortexing and incubation for 10 minutes on ice three times, lysates were centrifuged at max speed at 4°C for 20 minutes and supernatants were collected. Pierce BCA Protein Assay Kit (Thermo, 23225) was performed to determine the protein concentrations. 30 μg proteins were loaded to 8% acrylamide:bis-acrylamide gel. Gel was run at 90V for 30 min and at 120V for one and a half hours. Gel transfer was performed at 350mA for one and a half hours in a box full of ice. Non-fat dried milk powder was used for blocking the membrane for one hour at room temperature. KDM6A (CST 33510), HES1 (CST 11988), TLE1 (CST 4681S), b-actin (CST 3700) primary antibodies were used with 1/1000 dilution and CST 5151S, LICOR 926-68072 secondary antibodies were used with 1/30000 dilution. Nitrocellulose membrane was incubated overnight at 4°C with the KDM6A primary antibody; one and a half hours at room temperature with b-actin antibody. Secondary antibody incubations were done for one hour at room temperature. The membrane was imaged with the Li-COR ODYSSEY Clx machine in auto mode.

### 2.8. Identification of KDM6A mutation

DNA was extracted from cell lines BdEC, T24 and SV-HUC-1 using ZymoResearch Quick DNA Miniprep Plus kit (D4068). Mutation analysis was performed using the Archer VariantPlex Myeloid Panel (Diagnostica Longwood), an NGS panel containing the *KDM6A* gene. Mutation calling was performed using ArcherDx Analysis (Version 6.2.7) with default settings with Gen-ERA NGS service.

### 2.9. Plasmid Constructs and Transfection

To generate FLAG-KDM6A construct, pCMV_HA_UTX (Addgene, Plasmid #24168) vector was digested 2h at 37°C using Kpn1 and Not1 restriction enzymes, and KDM6A gene was removed from this vector. In order to add Flag-tag to KDM6A, 2-step PCR was set up using the Q5 polymerase (M0494S, Neb) with designed primers (see primer list, Table S3). PCR product was cut from the agarose gel and isolated with NucleoSpin^®^ Gel and PCR Clean-up kit (MN, 740609). Digested FLAG-tag KDM6A gene and digested pcDNA3.1(+)/myc-HisA (Invitrogen) vector were ligated 2h at room temperature using T4 Ligase (Neb, M0202S) in accordance with Neb Ligation Calculator (https://nebiocalculator.neb.com/#!/ligation). Ligation product (pcDNA3.1_FLAG_KDM6A) was transformed into E.coli 10beta bacterial strain. Growing colonies were isolated with the plasmid isolation kit (MN, 740588) and validation was done with Sanger sequencing. 80% confluent 10cm plate HEK293T cells were transfected with 7.5 μg plasmid DNA using Lipofectamine 3000 (Invitrogen, L3000015) reagent. Medium was changed 24h after transfection. To generate KDM6A truncated mutation like in T24 cell line, pcdna3.1_FLAG-KDM6A plasmid was digested 2h at 37 °C using Kpn1 and Not1 restriction enzymes, after ran on 1% agarose gel at 100V for 45 min, FLAG-KDM6A band was isolated with NucleoSpin Gel and PCR Clean-up kit (MN) from the gel. 2-step PCR was set up using the Q5 polymerase with designed primers for E895* mutation (see primer list, Table S3). After that, the same steps with generation of pcDNA3.1_FLAG_KDM6A plasmid were followed.

### 2.10. Co-IP

Proteins from untransfected and pcDNA3.1_FLAG_KDM6A or pcDNA3.1_FLAG_E895*mutant_KDM6A transfected HEK293T cells (48h post transfection) were isolated using 300 μl Lysis Buffer (50mM Tris-HCl pH:7.4, 150mM NaCl, 1mM EDTA, 1% TritonX-100, Protease Inhibitor Cocktail). For input controls 20 μl untransfected and transfected HEK293T cell lysates were used. Remaining proteins were bound overnight at 4°C on the rotator with Flag affinity gel (Sigma, A2220) in accordance with the manufacturer’s instructions. After the incubation, IP proteins were centrifuged and pellets were washed with TBS three times. Proteins for both input and IP fractions were boiled with SDS Sample Buffer (62.5nM Tris-HCl pH:6.8, 2%SDS, 10% (v/v) Glycerol and 0.002% Bromophenol Blue) at 95°C for 3 min and analyzed by western blotting.

#### Data analysis and structural modeling procedures

All the bioinformatics analysis was done in cooperation with IBG-BIP (our center’s bioinformatic platform).

### 2.11. Alignment and processing of next generation sequencing data

ChIP-seq data was aligned to the human genome (hg38) using nf-core/chip-seq pipeline (https://nf-co.re/chipseq/1.2.1). Using this pipeline, KDM6A ChIP peaks were called, peak annotation was done and KDM6A signals on peaks were quantified in CPM as well.

### 2.12. Peak calling and peak overlap

KDM6A peak calling was done using MACS2 (embedded in nf-core chip-seq pipeline) using the parameters: p<10^-5^ and FDR < 0.1 filter with narrow option. For these set of peaks called for T24, SV-HUC-1 and BdEC cell lines, the overlap among the peaks was checked using the “VennDiagram package” in R (Figure 1C).

**Figure 1.**
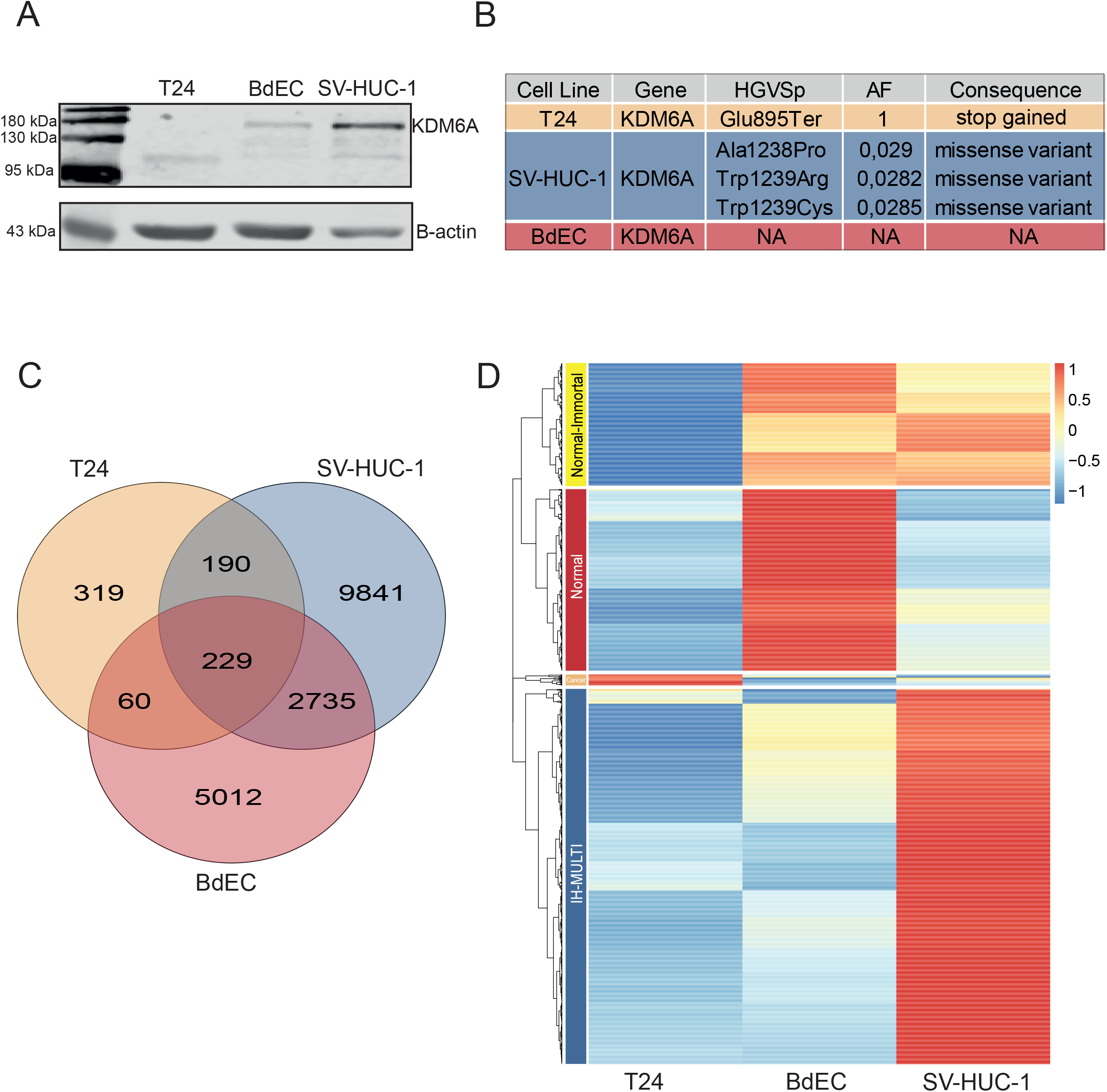
Occupancy and mutation profile of KDM6A in different bladder cell lines. (A) Western blot image displaying expression of KDM6A at protein level. (B) Image represents the details concerning depth, allele frequency and alternative alleles related to mutation status of KDM6A. (C) Venn diagram showing the overlap among KDM6A peaks called in T24, SV-HUC-1 and BdEC cell lines. (D) Heatmap shows the hierarchical clustering of KDM6A peaks, resulting in 4 clusters, Normal-Immortal (n=235), Normal (n=349), Cancer (n=21) and IH-MULTI (n=721).

### 2.13. Clustering of KDM6A peaks

Only the peaks which are located within ±2kb distance of transcriptional start site (TSS) of genes and fc value > 3 (value coming from MACS) were used for the clustering analysis. Before clustering, CPM counts are normalized for the library sizes and log2 of normalized values are used for plotting. Hierarchical clustering was done using “pheatmap package” (https://cran.r-project.org/web/packages/pheatmap/index.html) in R. Clusters were determined and visualized using the cutree=4 option in the package.

### 2.14. Analysis of T24 KDM6A peaks

TSS ±2kb filtered peaks in the Venn Diagram were used for the analysis. Peaks were ranked to their CPM normalized log2 values for each cluster. pheatmap R package was used for the visualization.

### 2.15. Transcription factor motif finding

Transcription factor motif finding for different KDM6A clusters was performed using findMotifsGenome.pl command of HOMER (http://homer.ucsd.edu/homer/motif/) with -size 300 parameter using genomic positions.

### 2.16. Gene ontology analysis

GO term analysis was done using ConsensusPathDB (http://cpdb.molgen.mpg.de/) with GO Level 3 and biological process (BP) options. Top 10 terms for ‘IH-MULTI’, ‘Normal’ and ‘Normal-Immortal’ clusters were identified and for those terms, for GO Level 3 were visualized using the dotplot function of the ggplot2 package (https://cran.r-project.org/web/packages/ggplot2/index.html). Only the terms with qval<0.01 filter were visualized.

### 2.17. Data visualization

ChIP-seq signal data visualization was performed using Bioconductor Gviz package (https://bioconductor.org/packages/release/bioc/html/Gviz.html). “BW” files were obtained from CPM normalized BAM and BAI files using deeptools “BamCoverage” functions’ default parameters. Visualization was performed with “BW” files using ‘horizon’ type of GViz package. GENCODE hg38 Comprehensive Gene Annotation Version 30 data was used as GeneRegionTrack for the annotation.

### 2.18. String protein interaction data visualization

For protein interaction data visualization Cytoscape (version 3.8.2) default style was used with String data.

### 2.19. HHEX and HES1 gene expression related associations

To obtain correlation heatmaps between expressions of HHEX and HES1, primary bladder cancer Hiseq data from TCGA BLCA 2017 (n=426) and bladder cancer cell lines RNAseq data (n=25) from CCLE (https://depmap.org/portal/download/all/ (CCLE_RNAseq_genes_rpkm_20180929.gct)) were used to generate the correlation heatmaps (Figure 5B, Figure S4A). Pearson correlation coefficients for the correlation of the expression of HHEX and HES1 and the genes associated with the term 'regulation of the developmental process' within the Normal-Immortal cluster (n=34) were calculated using RPKM normalized log2 expression values. ggplot2 (version 3.3.5) and pheatmap (version 1.10.12) packages in R were used for the visualization.

**Figure 2.**
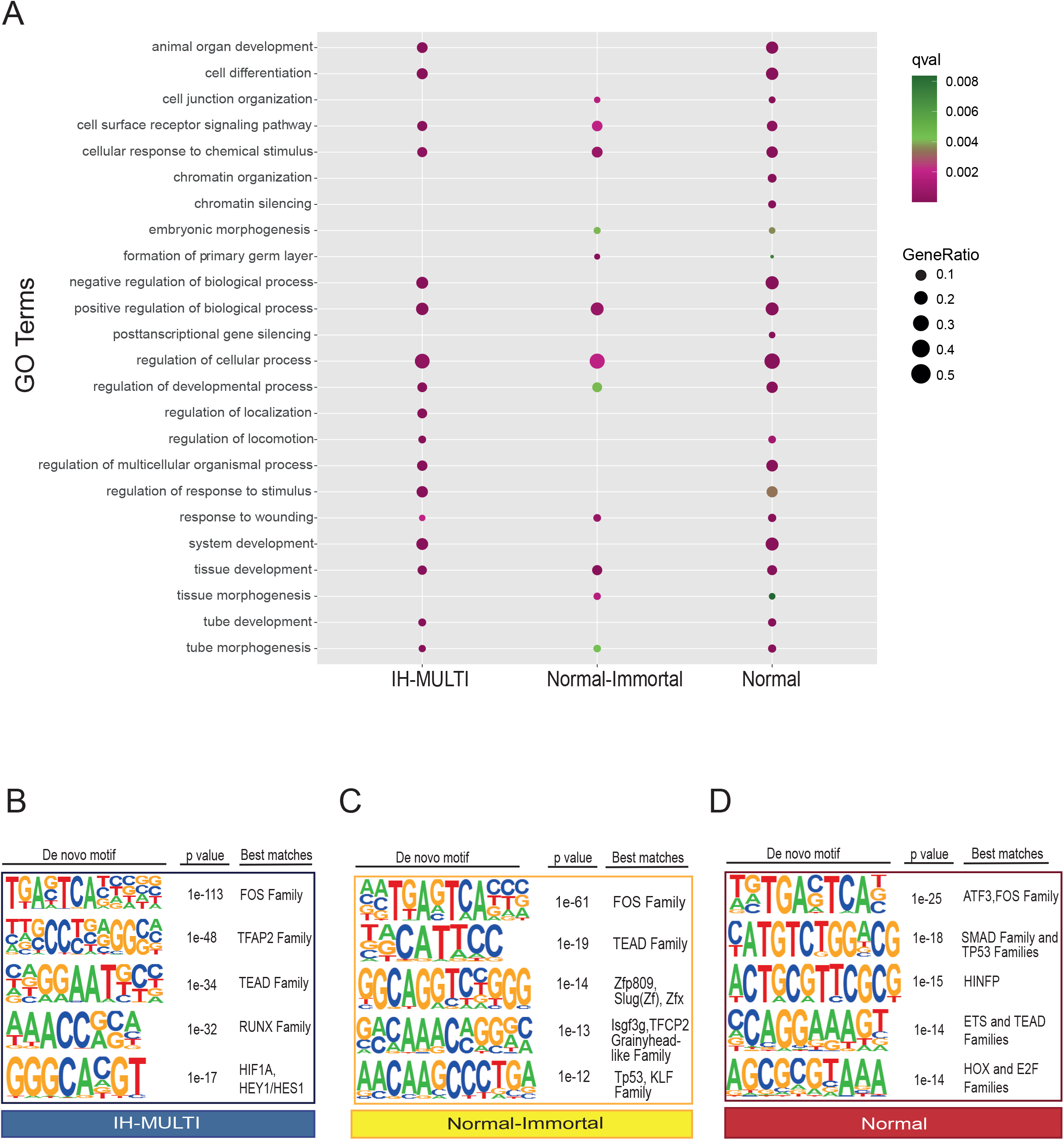
Functional analysis of differential KDM6A peak sets identified across the cell lines. (A) Dot plot image displaying the GO Term analysis performed for the genes associated with different clusters. (B-D) Transcription factor motif analysis performed for IH-MULTI (B), Normal-Immortal (C) and Normal (D) clusters. Up to 3 different transcription factor / transcription factor families are shown with the best motif matching scores.

**Figure 3.**
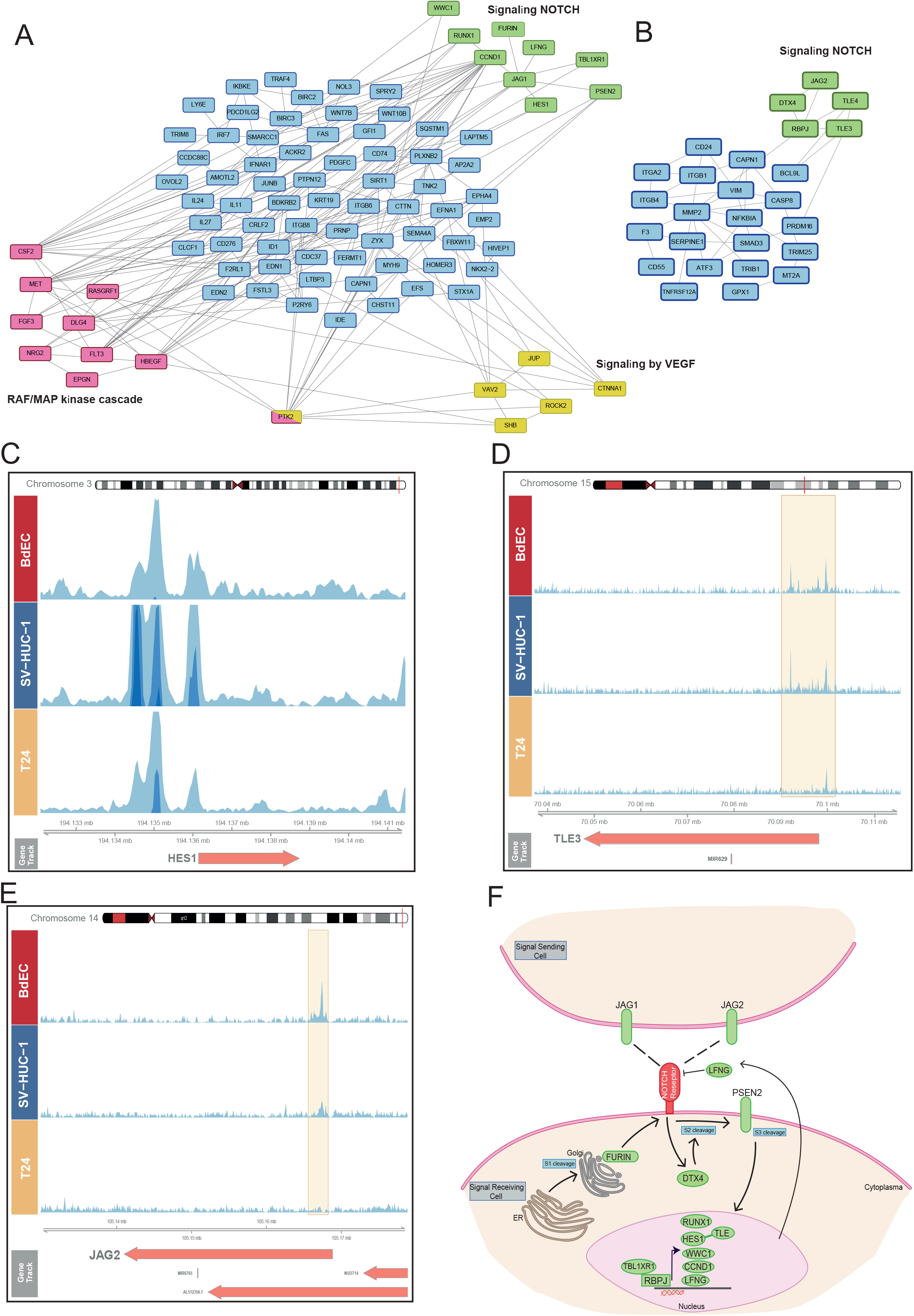
Genes regulated by KDM6A are implicated in notch signaling. (A-B) Analysis of the genes involved in cell surface receptor signaling for IH-MULTI (A) and Normal-Immortal clusters (B). Interaction data obtained from String was visualized using Cytoscape. (C-E) Snapshot images showing KDM6A signal in BdEC, SV-HUC-1, T24 cell lines at *HES1* (C), *TLE3* (D), and *JAG2* (E) loci. (F) Manually created cartoon showing the genes involved in Notch signaling and associated with KDM6A peaks (A, B). Genes marked with KDM6A signal are highlighted in green.

**Figure 4.**
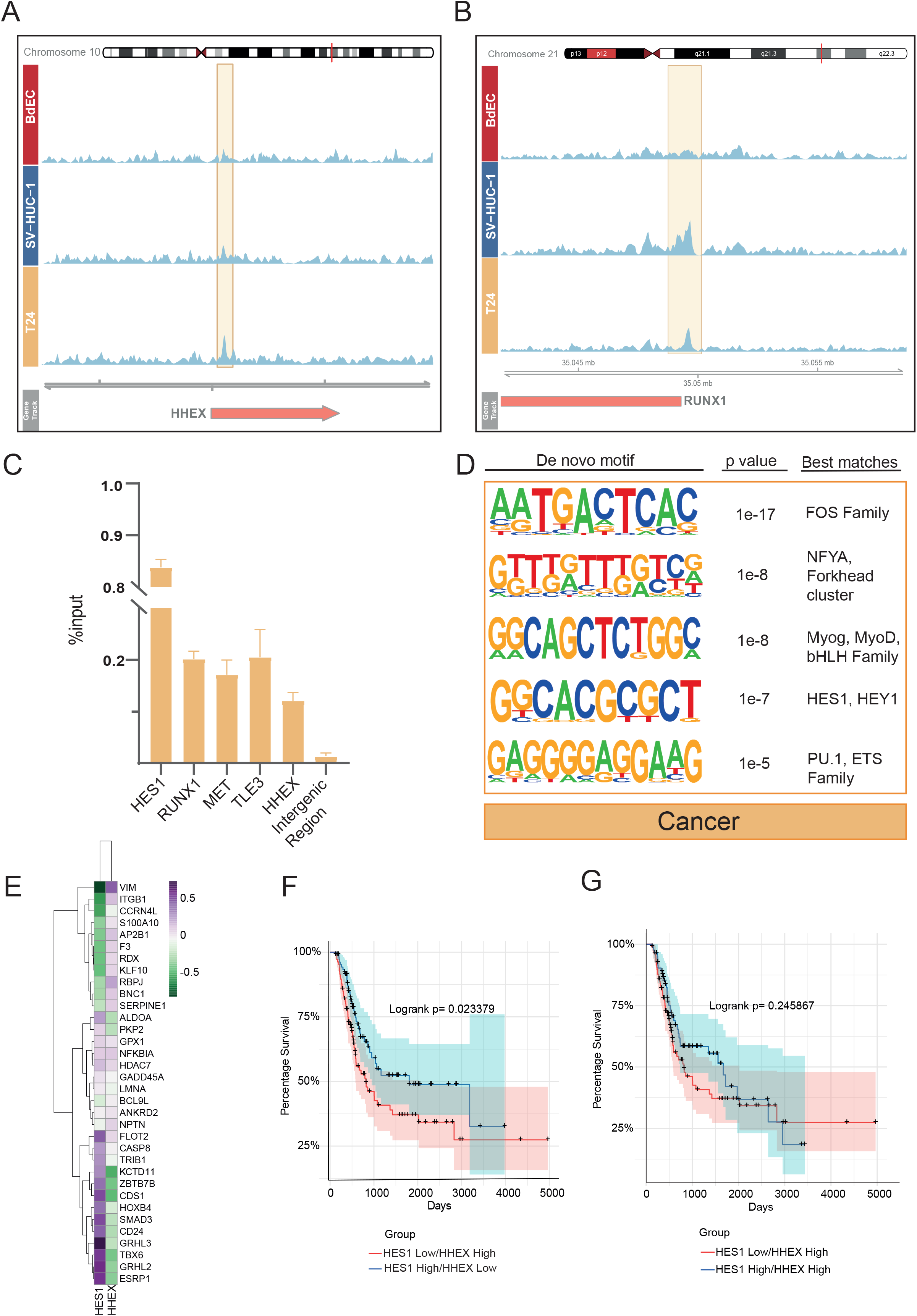
Genomic occupancy profile of truncated KDM6A in T24 bladder cancer cell line. (A-B) Snapshot images showing KDM6A signal in BdEC, SV-HUC-1, T24 cell lines at HHEX (A) and RUNX1 (B) loci. (C) ChIP-qPCR analysis of KDM6A enrichment in T24 cell line at selected loci. (D) Transcription factor motif enrichment analysis T24 KDM6A peaks linked with genes. (E) Heatmap demonstrates the correlation between the expression of HES1, HHEX and the genes involved in regulation of developmental process in primary bladder cancer. (F) Kaplan-Meier graph shows the overall survival of primary bladder cancer patients, grouped according to the expression status of HES1 and HHEX. (G) Kaplan-Meier graph show the overall survival of primary bladder cancer patients, grouped according to the expression status HES1 with ‘HHEX’ high criteria.

**Figure 5.**
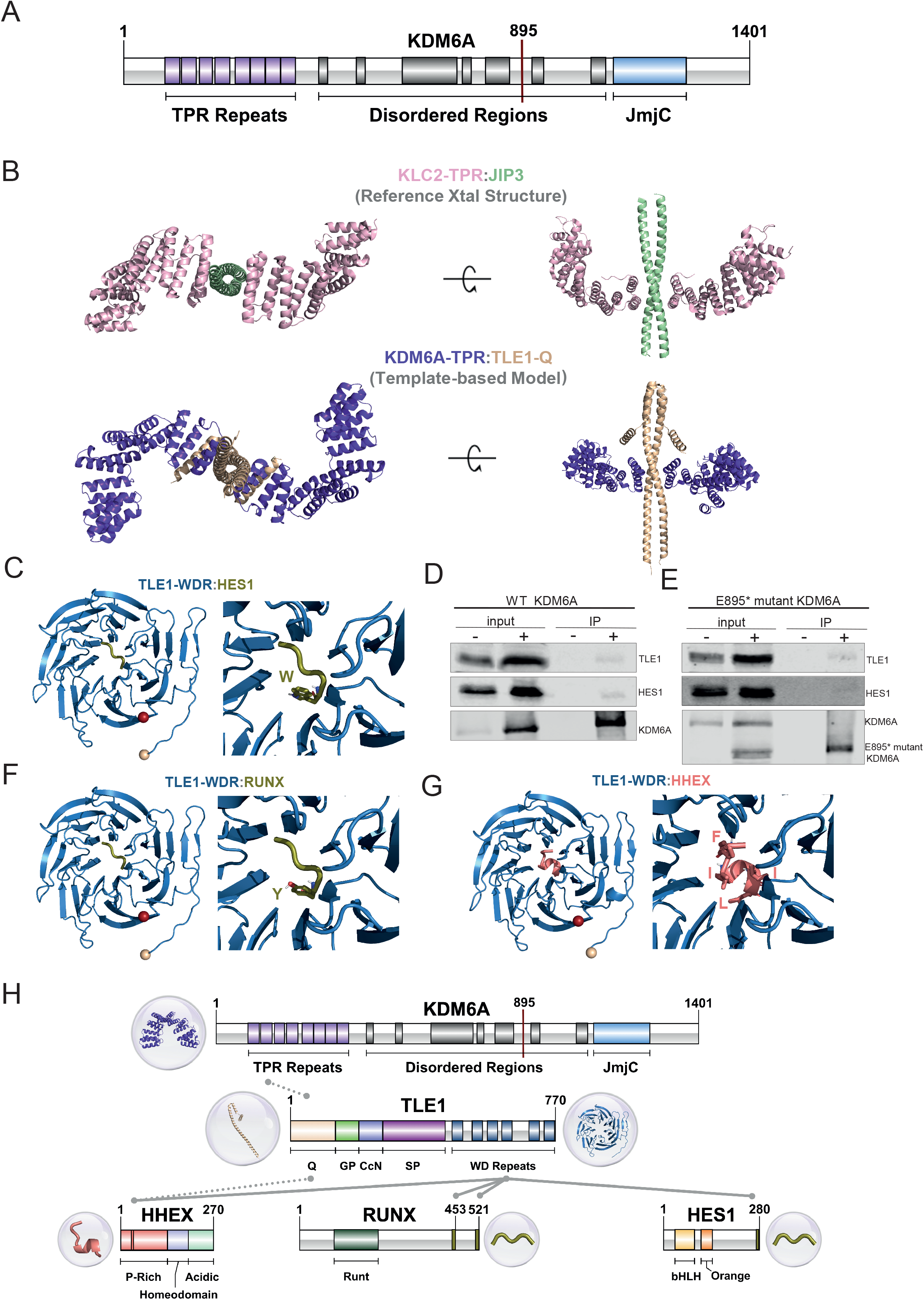
KDM6A TPR motif related interactions. (A) Domain illustration of KDM6A. (B) Modeling of KDM6A-TPR:TLE1-Q. The reference structure, containing KLC2-TPR (pink) and JIP3 (light green), is given in the first row (PDB ID: 6EJN). Our KDM6A-TPR (purple) and TLE1-Q (wheat) interaction model is shown in the second row. (C, F, G) Refined structures for TLE1-WDR containing interactions, together with the interacting motif closeups. N-ter (wheat), C-ter (red) represented with spheres. (D-E) Immunoprecipitation using anti-FLAG affinity gel in HEK293T cells expressing wild type flag-tagged KDM6A (D) and E895*mutant flag-tagged KDM6A (E). Western blot images show the bands detected for WT-KDM6A, truncated KDM6A, TLE1 and HES1. '-' and '+' denote the untransfected and transfect HEK293T cells, respectively for the input and IP samples. (H) Overall representation of interactions of co-repressors with KDM6A and TLE1 in domain level. Gray lines represent interacting regions between proteins, the source of the interaction is grouped as literature information and structural evidence, shown with dotted and solid lines, respectively. Three-dimensional structures are also provided for each interacting region.

### 2.20. Survival analysis

To generate Kaplan-Meier graphs showing patients survival times according to different expression levels of HHEX and HES1, gene expression RNAseq IlluminaHiSeq* (n=426) data and phenotype Curated survival data (n=436) belonging to TCGA Bladder Cancer (BLCA) cohort from Xena Browser Datasets [15] was used. Patients with survival data OS.time>120 (n=382) were included in Kaplan-Meier graphs. Low-high expression groups were formed by calculating the median values for HES1 and HHEX genes separately. Kaplan-Meier graphs were created with log-rank test according to OS.time values. Coin (https://cran.r-project.org/web/packages/coin/index.html) (version 1.4-2) and survival (https://cran.r-project.org/web/packages/survival/index.html) (version 3.2-7) R packages were used for the analysis.

### 2.21. Structural modeling

To support our *in vitro* findings, we modeled potentially interacting protein pairs linked to the truncated KDM6A interaction network, i.e., TLE1, HES1, RUNX1, RUNX2. For this, first, we retrieved the available relevant PDB structures from Protein Data Bank [16] We also gathered the structures of homologous protein complex systems that will help us model the unknown protein complexes. For the proteins with unknown structures, we used the deep learning-based protein modeling tool, AlphaFold2 (AF2) [17].

#### 2.21.1. KDM6A:TLE1 Interaction Modeling

KDM6A folds into tetratricopeptide repeat (TPR) and jumanji family (JmjC) domains. These two domains are connected by long-disordered regions between 437^th^ and 1079^th^ residues. Resolved KDM6A structures start from the 877^th^ residue, therefore they lack the N-terminal TPR domain (PDB IDs: 3AVR, 3AVS, 6FUK, 6FUL)[18, 19] The full-length TLE1 is 770 amino acids long protein, containing a glutamine-rich (Q), a glycine-proline-rich (GP), a nuclear localization signal containing central (CcN), a serine-proline-rich (SP), and a C-ter WD40 repeat domain (WDR). Among these domains, only Q and WDR domains are experimentally determined for the TLE1 protein (PDB IDs: 1GXR, 2CE8, 2CE9, 4OM2, 4OM3 and 5MWJ) [20–23] To model KDM6A:TLE1 complex, we focused on TPR and Q domains. TPR motif is formed by 34 amino acid long alpha-helix pair repeats and possesses a helical curvature. The structure of the Q domain is represented in 4OM2 and 4OM3 PDB entries [22]. In these structures, TLE1-Q forms a coiled-coil dimer through the interaction of 70 amino acid long alpha helices belonging to different TLE1s.

For the interaction modeling, initially we modeled the TPR domain with AF2 using default parameters [17]. Then, we modeled KDM6A-TPR:TLE1-Q with AlphaFold-multimer (data not shown here). Assessment of the AlphaFold models was carried out by analyzing: i) pLDDT scores showing the predicted confidence level of each residue, ii) predicted aligned error (PAE) graphs indicating predicted position errors of each residue, and iii) predicted TM (PTM) and interface predicted TM (iPTM) scores representing the overall accuracy of the complex and interface, respectively.

To generate good-quality models, we further searched the PDB Data Bank to find complexes containing TPR and coiled-coil interactions. 6EJN PDB structure was chosen as a reference here [24]. In this structure, the leucine zipper-like domain of JNK-interacting protein 3 (JIP3, a cargo protein) is found in a coiled-coil dimer form. The cargo interacts with two different mouse kinesin light chain 2 proteins (KLC2, a transport protein) through their first TPR motifs. The overall KLC2:JIP3 structure has a stoichiometry of 2:2. As this structure provides the binding pose for TPR and coiled-coil interaction, we generated our KDM6A-TPR:TLE1-Q model based on 6EJN complex. To build the targeted complex, we used the best-ranked KDM6A-TPR AF2 model. The coiled-coil TLE1 dimer is retrieved from the 4OM3 PDB entry [22]. We used the flexible structural alignment server, FATCAT, to superimpose the first three TPRs of KLC2 and KDM6A [25]. Obtained superimposed models were further used during structural fitting by PyMOL along with coiled-coil alignments, leading to our final model (The PyMOL Molecular Graphics System, Version 2.0 Schrödinger, LLC).

#### 2.21.2. TLE1:HES1 Interaction Modeling

HES1 contains basic-helix-loop-helix (bHLH) and orange domains and a Trp-Arg-Pro-Trp (WRPW) motif located at its C-terminal end. The WRPW motif is required for binding to Groucho/TLE family members. The PDB entry 2CE9 contains the interaction between the WRPW peptide and the Human Groucho/TLE WDR domain. In this structure, the WDR domain interacts with the WRPW peptide in a 2:1 fashion. This TLE1:HES1 complex is further refined with HADDOCK 2.4 webserver [26, 27]. HADDOCK is a user-friendly tool to perform biomolecular docking. After docking protocols, it also performs short molecular dynamics simulations to refine the interface by enabling sidechain movements. When turning off the docking procedures, HADDOCK completes the missing atoms in the input structures and improves the interface. To preserve the binding mode, we only applied the refinement step with unambiguous distance restraints to keep the peptide at its binding position (see advanced refinement protocol of HADDOCK 2.4 for details) [28].

#### 2.21.3. TLE1:RUNX1/2 Interaction Modeling

RUNX1 and RUNX2, hereinafter referred to as RUNX in the text, are characterized by approx. 120 amino acid long RUNT domain, required for DNA recognition. C-ter Trp-Arg-Pro-Tyr (WRPY) motif is known to interact with TLE1-WDR. We modeled this complex by taking the 2CE9 PDB as a template. During modeling, we mutated the last amino acid in the WRPW peptide to a tyrosine, to match the WRPY motif of RUNX and refined it with HADDOCK2.4, as described above.

#### 2.21.4. TLE1:HHEX Modeling

The primary structure of HHEX, also known as proline-rich homeodomain protein (PRH), consists of three domains: a N-terminal proline-rich (P-rich), a central homeodomain interacting with DNA, and a C-terminal acidic domain regulating transcription activity. 137 amino acid long P-rich domain includes an EH1 motif (FxIxxIL) providing an interaction surface for TLE1. From the Protein Data Bank, we retrieved the 2CE8 structure, showing the interaction of TLE1-WDR with EH1 peptide (FSIDNIL) from human Goosecoid (GSC), another homeobox protein. To mimic the sequence of the EH1 motif in HHEX, FYIEDIL, we mutated the mismatched amino acids. Mutation and refinement procedures were performed as outlined above.

Further, we used AlphaFold-multimer to model the interaction between TLE1-Q and HHEX [29]. AlphaFold-multimer allows the prediction of multimeric protein complexes with a known stoichiometry. Here, our input sequences contain the N-terminal 131 and 98 amino acids of TLE1 and HHEX, respectively. We modeled the TLE1:HHEX complex with a stoichiometry of 2:1. Default parameters were used for multimer modeling in our local machine. Assessment of the models was performed as described above.

### 2.22. Statistical analysis

R software was used for all data analysis and plots. Kaplan-Meier graphs were created using logrank test according to OS.time values. ChIP-qPCR and gene expression data were visualized using GraphPad Prism 9. Pearson correlation coefficients were calculated for the correlation between gene expression values.

## 3. Results

### 3.1. KDM6A mutation status, identification, and classification of KDM6A peaks in bladder cell lines

We performed KDM6A ChIP-seq for three different cell lines, BdEC (normal primary bladder epithelial cells), SV-HUC-1 (immortalized bladder epithelial cells), and T24 (muscle invasive bladder cancer cell line). Among the cell lines analyzed, T24 had a known stop-gain mutation (position: E895). Prior studies showed that this mutation was homozygous, and western blot analysis revealed a protein product about the size of 110 kDa [30]. Our own Western blot analysis also confirmed the presence of KDM6A in all three cell lines, and a truncated mutated product for T24 cell line (Figure 1A). The mutant KDM6A observed for T24 cell line lacks the histone demethylase domain. To be complete about the mutation status of the cell lines used in the study, we determined the mutation status of KDM6A in all three cell lines ourselves as well. As expected, T24 cell line has a homozygous truncating mutation at position 895 for KDM6A. SV-HUC-1 cell line had missense mutations at positions 1238 and 1239 with allele frequency of 0.029 and 0.0282, while BdEC was wild type for KDM6A (Figure 1B). We considered the missense mutation observed for SV-HUC-1 as a reflection of clonal mosaic pattern and to have a negligible effect on chromatin binding.

We called KDM6A peaks in the three cell lines from our ChIP-seq data, resulting in 8050, 13039, 819 number of peaks in BdEC, SV-HUC-1 and T24 cell lines, respectively. 16% of peaks were shared between BdEC and SV-HUC-1 cell lines, whereas the total number of peaks called for T24 cell lines were rather limited. Here, only 229 number of peaks were common among the all three cell lines used (Figure 1C). We then proceeded with the identification of KDM6A peaks with differential enrichment of the KDM6A signal. This analysis resulted in four distinct clusters, which we named as ‘normal-immortalized shared’ (Normal-Immortal) (n=235), ‘Normal’ (n=349), ‘Cancer’ (n=21) and ‘immortalized high-multiform in cancer and normal’ (IH-MULTI) (n=721) (Figure 1D).

### 3.2. Characterization of genomic loci with differential KDM6A signal

After determining the genomic regions with differential KDM6A signal in the cell lines analyzed, we annotated them with functional features. First, we identified the genes associated with the peaks located in the clusters we defined (Figure 1D) (peaks located within ± 2kb distance of transcriptional start sites (TSS), see Methods) and performed gene ontology analysis for the genes identified in different clusters. Genes associated with the ‘Cancer’ cluster were not significantly enriched for a gene ontology term. For the other three clusters, terms related to general developmental functions showed similar enrichment patterns for all three clusters. ‘Normal’ and ‘Normal-Immortal’ clusters were similarly enriched for ‘cell junction organization’ (Figure 2A, Table S1). On the other hand, we identified ‘chromatin organization’ related terms to be significantly enriched only in ‘Normal’ cluster (Figure 2A). These genes exceptionally mainly consisted of the replication-dependent histone genes (Figure S1A–S1B). In contrast, non-canonical histones did not have KDM6A signal in any of the three cell lines (Figure S1C–S1D).

Next, we performed transcription factor motif analysis for the peaks belonging to different clusters (Figure 1C). All clusters were similarly enriched for FOS and TEAD family transcription factor motifs (Figure 2B–2D). We identified enrichment of several members of Zinc finger family, Grainyhead-like family, and p53 transcription factor motifs at peaks located within ‘Normal-Immortal’ cluster, while RUNX and bHLH family of transcription factors (HIF1A/HEY1) were enriched at peaks within ‘IH-MULTI’ cluster (Figure 2B–2C). ‘Normal’ cluster showed the enrichment for Smad family, p53 and HINFP, a transcription factor involved in transcriptional regulation of histone H4 [31] (Figure 2D). As the identification of transcription motifs only does not directly provide information about the functional involvement of these factors, we checked the expression of several transcription factors with motifs enriched in these clusters, which might be relevant for bladder cancer biology. Combined with the motif finding results, these data (Figure S2) suggested that KDM6A might potentially cooperate with TP53 family, GRHL2 and FOSL2 transcription factors to drive gene expression associated with epithelial characteristics while coordination with transcription factors such as FOSL1, previously described to be involved in invasive character of bladder cancer [32, 33], might contribute to the gene expression programs driving tumorigenesis.

### 3.3. KDM6A occupies the genes involved in Notch signaling

To decipher the signaling pathways associated with KDM6A function, we analyzed the term ‘cell surface receptor signaling pathway’, which are similarly enriched for different clusters. ‘IH-MULTI’ class showed enrichment for Raf/MAP kinase cascade and VEGF signaling (Figure 3A). Surprisingly, genes involved in cell surface receptor signaling showed clustering of the genes associated with Notch signaling for both ‘IH-MULTI’ and ‘Normal-Immortal’ clusters (Figure 3A–3B). ‘Normal’ class also had enrichment for Notch signaling. However, we discovered that those mostly consist of histone genes, reflecting the enrichment of the genes involved in chromatin organization in ‘Normal’ cluster (Figure S3). Notch signaling has a key role for normal urothelium development and its deregulation is implicated in bladder tumorigenesis [34, 35]. We identified KDM6A signal at the promoter regions of several prominent Notch signaling related genes, such as *HES1*, transcriptional repressor, and *TLE3/4*, transcriptional co-repressor, both critical for cell fate decisions [36], *JAG1* and *JAG2*, ligands of the NOTCH receptor genes [37], *RBPJ*, a DNA binding protein which can complex with cleaved Notch to activate transcription [38] in all three cell lines with different signal intensities (Figure 3C–3E, Figure S4A), covering all prominent components of Notch signaling (Figure 3F). Upon checking the expression of KDM6A-occupied Notch signaling genes in the analyzed cell lines, we identified that occupancy level of KDM6A correlates with the expression level of the genes overall (Figure S4B), supporting the relationship between active gene expression and chromatin organization by KDM6A. However, it should be noted that relative levels of transcription factors and repressive chromatin remodeler complexes might be critical in regulation of gene expression by KDM6A as it is shown in literature before [10, 39]. Collectively, our results underscore the role of KDM6A in regulation of Notch signaling and its potential deregulation in bladder cancer.

### 3.4. Truncated KDM6A associates with genes deregulated in cancer

We were able to find only a limited number of peaks annotated with a gene for the ‘Cancer’ cluster (n=20) (Figure 1D). Performing a gene ontology analysis for this set of genes did not result in any significant GO terms. However, as we have shown that the truncating mutation at position 895 results in a protein product and this product still shows some chromatin binding profiles. Thus we re-inspected all T24 KDM6A peaks within ±2kb distance to TSS (Figure 1C) independent of their cell line specificity (n=50) (Table S2) to decipher more mechanistic insights which might be associated with truncated KDM6A. In relation to our initial clustering analysis (Figure 1D), all T24 KDM6A peaks were mostly either located in ‘IH-MULTI’ (%42) or ‘Cancer’ (40%) clusters. The peaks with the highest signal intensity are also found in these respective clusters, too (Figure S5A). Among the genes occupied by KDM6A in T24 cell line were Notch signaling family members, such as *HES1* and *TLE3* as we have already shown in Figure 3 and genes deregulated in cancer such as *RUNX1, HHEX*, and *MET* [40–42] (Figure 4A–4B, Table S2). Additionally, ChIP-qPCR validation showed the occupancy of mutant KDM6A at these loci, confirming the presence of truncated KDM6A at these genomic regions, rather than being an artifact (Figure 4C).

As the bladder cancer cell line included in our study, T24 is not wild type for *KDM6A*, we additionally compared our data to a published bladder cancer cell line KDM6A ChIP-seq data. The used data is for wild type KDM6A expressing-UMUC-1 luminal bladder cancer cell line, where the authors performed ectopic expression of wild type KDM6A on UMUC-1 cells, which is normally *KDM6A* mutant and lacks KDM6A expression [43]. This comparison revealed that overall 114 out of 798 KDM6A peaks identified for T24 cell line overlapped KDM6A peaks called for wild type KDM6A expressing-UMUC-1 cells (Figure S6A). The low rate of intersection might be either attributed to the wild type vs truncated KDM6A function or different character of these two cancer cell lines. However, analysis of genes associated with overlapping peaks pointed the occupancy of KDM6A in wild type KDM6A expressing-UMUC-1 cells at two key Notch signaling genes, *HES1*, and *TLE3*, as well (Figure S6B).

After performing a transcription factor motif analysis at T24 KDM6A peaks linked with genes, similar to other clusters, we identified FOS family members at the top. We also found HES1 and other bHLH family motifs as enriched (Figure 4D). This result suggested that KDM6A is involved in transcriptional regulation of HES1, and in turn KDM6A and HES1 interaction might be implicated in regulation of genes in bladder cancer. Although we determined HES1 motif to be enriched primarily in KDM6A peak regions identified in T24 cell line, our transcription finding motif analysis on IH-MULTI cluster showed the potential enrichment of HEY/HES1 family for this cluster as well. Additionally, although ranking low, we identified HES1 motif to be enriched at Normal-Immortal cluster (data not shown). Together, these data suggest that HES/HEY transcription factors are critical players of KDM6A dependent gene regulation with varying degrees.

Among the genes occupied by truncated KDM6A in T24 cell line, HHEX especially gained our attention. Like HES1, HHEX is also a transcriptional repressor involved in many developmental processes [44]. HHEX has the highest expression pattern in T24 cell line among the cell lines analyzed in contrast to the lowest expression of HES1 in the same cell line (Figure S5B). We wondered whether these two KDM6A-occupied repressors, *HHEX* and *HES1*, act differentially on the regulation of genes critical for urothelium development. Therefore, we plotted the correlation between HHEX, HES1 expression and the expression of the genes associated with the term ‘regulation of developmental process’ for Normal-Immortal cluster (Figure 1D) for primary bladder cancer [45–48]. This analysis revealed a striking positive and corresponding negative correlation for the expression of developmental transcription factors such as *GRHL2, GRHL3, ZBTB7B, TBX6* [49–51], and the expression of HES1 and HHEX, respectively (Figure 4E). Making a similar analysis using CCLE bladder cancer cell line data showed comparable patterns for these factors albeit with a bit more scattered clustering (Figure S5C). Based on this contrasting association observed for HES1 and HHEX, we next classified the MIBC patients (TCGA, 2017) according to the expression status of HES1 and HHEX individually and checked the survival. This analysis revealed a statistically significant better overall survival for the patients expressing high levels of HES1 (Figure S5D), while the patients expressing high levels of HHEX showed worse prognosis without statistical significance (Figure S5E). We further classified the patients according to the co-expression status of HHEX and HES1 as ‘HES1-high/HHEX-low’ and ‘HES1-low/HHEX-high’. Our analysis showed that patients, which are in the ‘HES1-high/HHEX-low’ group showed significantly better prognosis compared to the patients belonging to the ‘HES1-low/HHEX-high’ group, especially providing an advantage for midterm survival (1000-3000 days) (Figure 4F). On the other hand, with a high HHEX expression level, high HES1 expression does not bring an advantage for survival compared to low HES1 expression (Figure 4G). These results suggest that although HES1 expression level seems to be the main player in HES1-HHEX interplay with regard to the patient survival, higher expression level of HHEX disrupt the beneficial effects of high HES1 expression.

### 3.5. Structural modeling and co-IP illustrate previously uncovered KDM6A-TLE-HES1 interactions

We identified that truncating mutation at position 895 of KDM6A still results in a protein product and is associated with chromatin. We also saw that the truncated KDM6A might still interact with the transcription factors, where HES1 was determined as one prominent candidate. Therefore, we concentrated on understanding the effect of KDM6A truncation on its chromatin and transcription factor binding landscape.

The full-length KDM6A is 1401 amino acids long, containing eight TPR motifs at its N-terminal and a JmjC domain at its C-terminal region. The function of KDM6A-TPR is yet to be resolved. The JmjC domain, on the other hand, was demonstrated to regulate KDM6A’s enzymatic activity for demethylating H3K27 [52]. The TPR and JmjC are connected through an approx. 600 amino acids long intrinsically disordered region (IDR) (Figure 5A). There is no structural information available on the full-length KDM6A, except its AlphaFold 2 (AF2) model deposited at the EBI’s database (Figure S7A) [53]. The prediction quality of this model shows that AF2 could predict the fold of TPR and JmjC domains accurately, whereas it could not predict the organization of the IDR and the orientation of the N- and C-terminal domains with respect to each other (Figure S7B).

Upon investigating the interacting partners of KDM6A in BioGRID database (https://thebiogrid.org/), we realized that one interacting partner is TLE1. TLE1, belonging to the Groucho/Transducin-Like Enhancer of Split (TLE) family, is a co-repressor involved in diverse developmental functions [54]. Further, it is also known that TLEs can interact with HES1 [54]. Previously, it was shown that first three TPRs of KDM6A interacts with the N-terminal Q domain of TLE1 [55]. Expanding on this literature information, we explored the means of KDM6A:TLE1 interaction.

TLE family co-repressors (TLE1-4) all have a conserved N-terminal Q domain and C-terminal WDR domain, which are structurally resolved as isolated domains (Figure S8) [54]. From these structures, we know that Q domain is responsible for the tetramerization of TLEs by forming parallel coiled-coil dimers (Figure S8, [22, 56]) To model KDM6A-TPR:TLE1-Q interaction, we looked for homologous systems, though we could not obtain any structure with meaningful sequence homology. Therefore, we checked all the available TPR structures that bind to a coiled-coil. As a result, we found the PDB entry 6EJN, a mouse KLC2 TPR domain binding to the JIP3 leucine zipper domain [24]. In this structure, KLC2-TPR carries its cargo JIP3 in a 2:2 fashion, where two independent TPRs bind to the coiled coil dimer of JIP3 through their first TPR (Figure 5B). As this TPR interaction information overlaps with the one of KDM6A:TLE1, we decided to use 6EJN as a template for our KDM6A-TPR:TLE1-Q complex modeling. Although there are no significant sequence similarities between KDM6A-TPR:TLE1-Q interaction system and KLC2-TPR:JIP3, upon one-to-one structural alignment (with FATCAT [25]), we could build a working structural model for the KDM6A-TPR:TLE1-Q complex (Figure 5B). This model portrays the first structural hypothesis on how KDM6A can carry out its non-enzymatic tasks in the absence of its JmjC domain. It also suggests that KDM6A-TPR:TLE1-Q complex is formed in a 2:2 fashion, which is prone to further experimental validation.

We also know that TLE1 interacts through its WDR domain with HES1’s C-terminal WRPW motif (2CE9) [21] (Figure 5C). This tells us that TLE1 interacts with KDM6A and HES1 via its different domains. By this, we can suggest that KDM6A, TLE1, and HES1 form a complex functioning together. To probe such an interaction, we overexpressed FLAG-tagged full-length KDM6A and performed a co-immunoprecipitation experiment. The antibody used for the detection of TLE1 recognizes the other TLE proteins (TLE2, TLE3, TLE4) as well based on the conservation of all functional domains in these proteins [57]. However, for simplicity, we refer to the detected protein as ‘TLE1’. Our co-immunoprecipitation results demonstrated the potential presence of both TLE1 and HES1 in the same complex with KDM6A (Figure 5D). As this experiment was performed with wild type KDM6A having all functional domains, to further illustrate that TLE1:KDM6A interaction occurs via TPR domain of KDM6A, we carried out another co-immunoprecipitation experiment using FLAG-tagged truncated KDM6A, mimicking the E895* mutation observed in T24 cell line. These results also revealed the association of truncated KDM6A with TLE1, and HES1 as well, albeit a bit weaker compared to full-length KDM6A (Figure 5E). Collectively, these data emphasize the role of TPR domain in these protein interactions. Overall, KDM6A, TLE1 and HES1 interaction might be puzzling, as TLEs and HES1 are involved in transcriptional repression and KDM6A is responsible for active chromatin organization. Several studies have shown that TLE/HES1-mediated repression can be resolved by the action of transcription factors, turning the repressive complex into an activator. For instance, RUNX2 has been determined to interact with the TLE-WDR domain and thus can interrupt the interaction of the TLE-WDR domain with HES1 and resulting in transcriptional derepression [58]. RUNX proteins are 453 and 521 amino acid long, containing WRPY (WRPW-like) motifs in their C-termini. So, RUNX’s WRPY might compete with WRPW of HES1 for the same binding site on TLE1 (Figure 5F). Another study also showed that HIPK2 can interact with TLE1/HES1 complex and mediate transcriptional activation critical for cortical neurogenesis [59]. Indeed, our model for the interaction of RUNX family transcription factors, enriched at KDM6A peaks belonging to IH-MULTI cluster (Figure 2B), with TLE1 illustrated this association (Figure 5F). Furthermore, additional repressors can interact with TLE:HHEX transcriptional repressor, another KDM6A-regulated gene we identified (Figure 4A). HHEX consists of P-rich, homeo, and acidic domains.

It has a FxIxxIL motif residing in between 32 and 38 residues in the P-rich domain. HHEX has been suggested to interact with both TLE1-WDR and TLE1-Q via its N-terminal 98 residues [60]. The studies narrowed down the interface of HHEX with TLE1-WDR to FxIxxIL motif in the N-terminal HHEX as shown in our structural model [61] (Figure 5G). Though the N-terminal HHEX has been suggested to interact with TLE1-Q, we could not obtain a confident model showing this interaction. Therefore, our AF2 model is presented in SI. (Figure S9).

Collectively, all these results and the knowledge extend the interaction network of TLE1, suggesting a dynamic interaction network depending on the domain characteristics of the interactors (Figure 5H). Based on this, we propose a regulatory mechanism where KDM6A associates with TLE co-repressor complexes, and upon action of a transcriptional activator, KDM6A exerts its role in active chromatin organization and transcriptional activation.

## 4. Discussion

*KDM6A* is one of the most frequently mutated genes in bladder cancer [2]. Therefore, understanding the function of KDM6A in normal and tumorigenic bladder is essential. In this study, we performed KDM6A ChIP-seq in normal, immortalized and tumorigenic bladder cell lines. Our results showed differential KDM6A occupancy across the cell lines analyzed.

Among the KDM6A groups we defined, ‘Normal’ cluster was especially associated with the genes involved in chromatin organization, mainly consisting of replication dependent histone genes. We identified ‘Normal-Immortal’ and ‘IH-MULTI’ cluster to be enriched for the genes involved in Notch signaling, including the ligands, repressor, co-repressor proteins critical for this pathway. Notch signaling is implicated in normal urothelium development [35] and is known to be deregulated in bladder cancer [62]. Our findings here, which shows the relationship of KDM6A with Notch signaling provides additional insights about what goes wrong in the cancer samples when KDM6A is mutated. Previously, association of KDM6A with Notch signaling has been shown for neural crest cells in connection to Kabuki syndrome [63], reprogramming of germ line cells to neurons [64] as well.

The T24 cancer cell line we used in this study has a homozygous truncating mutation at position 895 for KDM6A. Overall, we identified much lower number of KDM6A peaks for this cell line (Figure 1C), still we realized that some loci showed strong KDM6A occupancy rather than being an artifact. This finding highly suggested that truncated KDM6A still preserves some of its chromatin binding abilities. Thus, we inspected this mutant product in more detail in terms of cooperation with transcription factors and structural modeling as well. Our transcription factor motif analysis implicated HES/HEY family to be associated with truncated KDM6A in T24 cell line. KDM6A has TPR motifs located at the N-terminus and a demethylase domain at the C-terminus. Our modeling showed that truncated KDM6A in T24 has the TPR motifs, which can fulfill the protein-protein interactions. In fact, using co-immunoprecipitation experiments, we show that KDM6A, TLE1, and HES1 are in the same complex. However, it should be noted that immunoprecipitation does not provide a complete proof for direct interaction of these proteins. We propose that TPR domain of KDM6A is in interaction with repressor regulatory factors, which is also in interaction with transcriptional activators. Upon receiving the correct signal, KDM6A loses its interaction with repressor regulatory proteins and activates transcription potentially via its JmjC demethylase domain. In the case of truncated KDM6A, TPR domain might interact with repressors and co-repressors such as HES1/TLE1. Also, the lack of JmjC domain may result in repression of the key differentiation genes while some oncogenic ones might be activated with the cooperation with transcription factors implicated in tumorigenesis.

HES1 is normally required for the timely differentiation of many cell types. It is known that function of HES1 in cancer might be abrogated, and high HES1 expression might be associated with less differentiated tumors [36]. Recently, a study investigating the expression of the genes involved in Notch signaling in normal bladder, papillary and non-papillary bladder tumors showed that overall the expression of HES1 is higher in both papillary and non-papillary tumors compared to normal, while its expression is higher in papillary tumors in comparison to non-papillary tumors [65]. Thus, we suggest that depending on different mutations observed for KDM6A, and the presence of different activator proteins, KDM6A-TLE1-HES1 regulatory axis can be manipulated in diverse ways boosting the tumorigenesis.

Our results further show that the interactions of TLE1 and subsequently with KDM6A might be quite dynamic, where associations and dissociations can be established depending on the domain organization and the relative amount of interacting proteins. In KDM6A-TLE1-HES1 regulatory axis, we suggest that another repressor, HHEX, which is highly expressed in T24 cell line comes into this regulatory scheme via its interaction with TLE1. The literature about the role of HHEX in bladder cancer is very limited. One study suggested that expression level of HHEX in superficial bladder cancer is lower compared to normal bladder [66]. Here, our results based on MIBC data (TCGA, 2017) suggest that high expression of HHEX shows a more oncogenic behavior and is associated with worse survival. Checking the results of the published study actually shows that compared to superficial bladder cancer, invasive cancer shows slightly higher level of HHEX expression [66]. Together, existing literature and our data propose a model that expression level of HHEX might show differential effects depending on the interplay with other repressors such as HES1 and the status and the grade of bladder tumors.

KDM6A has a JmjC demethylase domain functioning in demethylation of H3K27 [18, 52]. However, so far multiple studies showed that function of KDM6A in differentiation and development is largely independent of its demethylase activity [52, 63]. A recent study also demonstrated that expression of wild type KDM6A or KDM6A-demethylase-mutant in a KDM6A-null bladder cancer cell line resulted in similar KDM6A genomic localizations [43]. It was additionally shown that IDR region of KDM6A is critical for the creation of KDM6A liquid condensates and functioning of MLL4 [67]. Existing literature and our findings highly suggest that TPR motifs present at the N-terminus of the KDM6A is very critical in terms of specification of the interaction partners of KDM6A.

Collectively, our results provide many critical aspects about the distinct KDM6A occupancy in bladder cell lines with different characteristics and the potential interacting transcription factors. Importantly, we believe that KDM6A-TLE1-HES1 regulatory axis we defined makes a valuable contribution to the understanding of epigenetic deregulation in bladder cancer with regards to the mutations observed in KDM6A.

## Author Contributions

Conceptualization, G.Ö.Y., S.S., E.K., G.K., and S.E.O.; methodology, G.Ö.Y., B.S., A.B.; software, validation, formal analysis, investigation, data curation, visualization G.Ö.Y, B.S., A.B., A.E., A.A.; writing, review and editing, G.Ö.Y., B.S., S.S., E.K., G.K., and S.E.O.; supervision, project administration, funding acquisition, S.E.O. All authors have read and agreed to the published version of the manuscript.

## Supplementary Information for

**Table S1.** Results of Gene Ontology Analysis performed for the clusters in Figure 1D.

**Table S2.** Genes associated with all T24 KDM6A peaks within ±2kb distance to TSS.

**Table S3.** List of primers used in the study.

**Figure S1.**
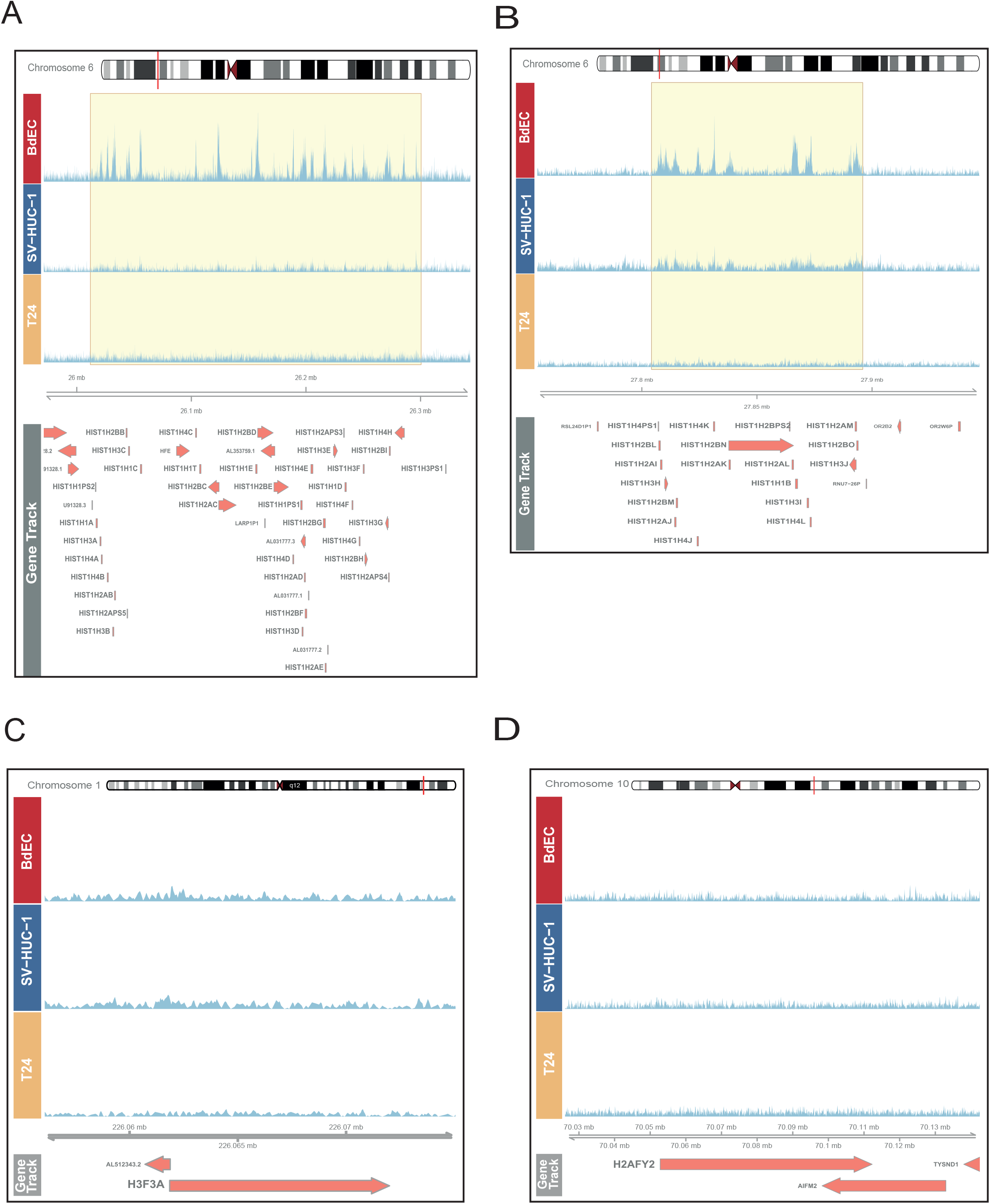
KDM6A localizes to the replication-dependent histone genes in normal bladder cell line. Snapshot images show the occupancy of KDM6A at canonical histone genes, *HIST1* (A-B), and non-canonical histone genes, *H3F3A* (C) and *H2AFY2* (D).

**Figure S2:**
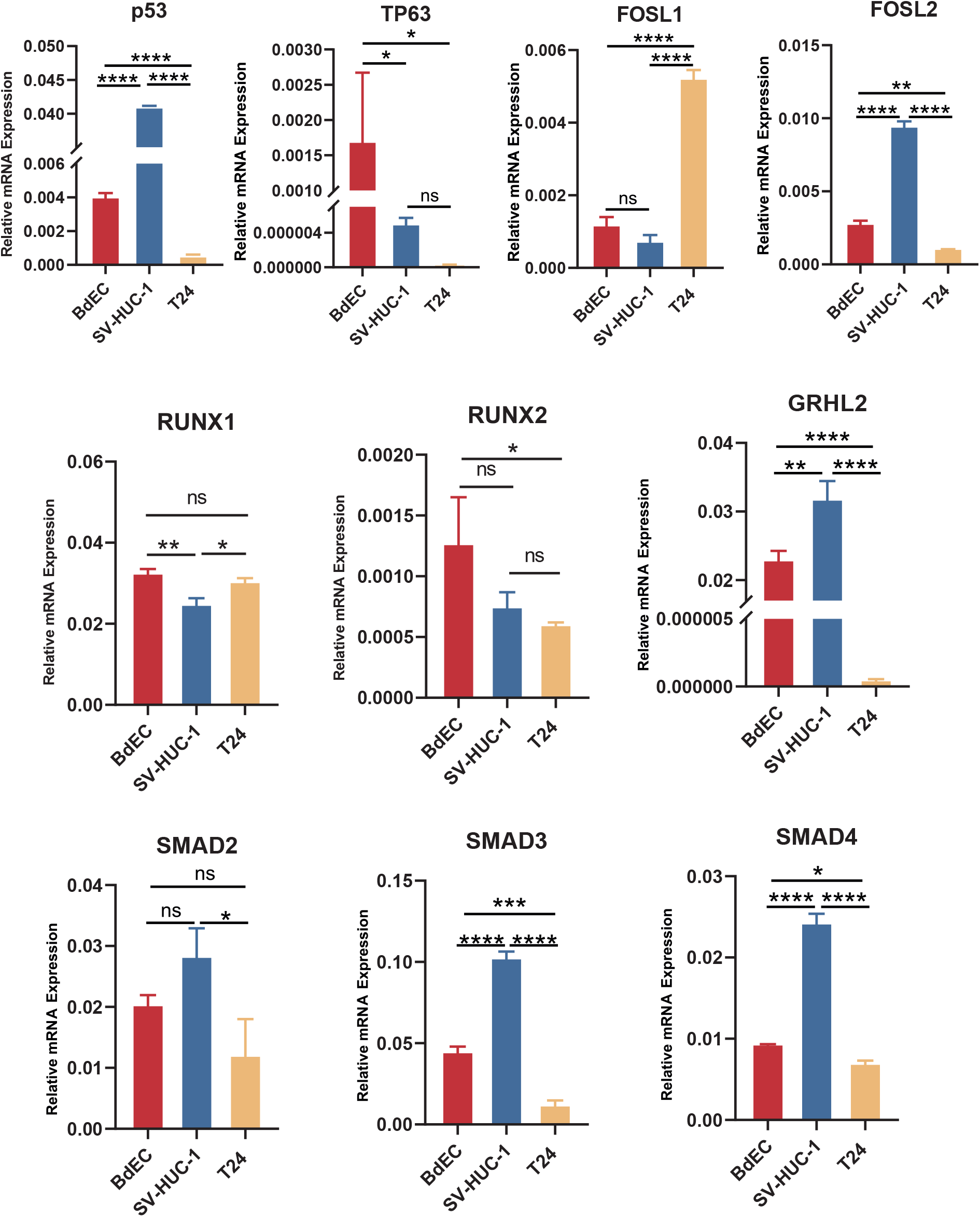
Expression of transcription factors in BdEC, SV-HUC-1 and T24 cell lines. Barplots show the relative expression level of *P53, TP63, FOSL1, FOSL2, RUNX1, RUNX2, GRHL2, SMAD2, SMAD3, SMAD4* as determ *i* ned by RT-qPCR analysis. Error bars depict the sd of three technical replicates. * pvalue < 0.05, ** pvalue < 0.005, *** pvalue < 0.0005, **** pvalue < 0.0001, ns = not significant

**Figure S3.**
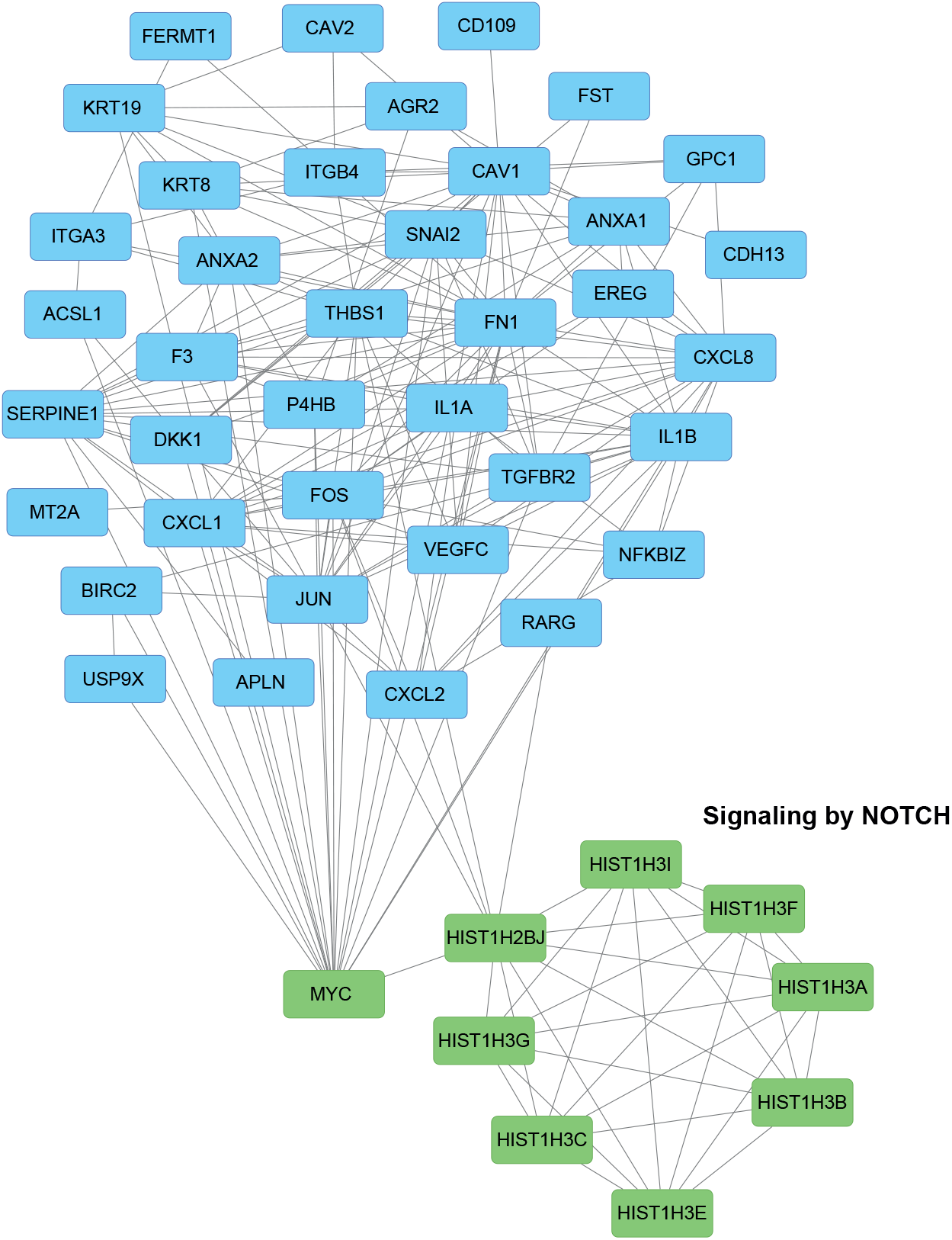
Analysis of the genes involved in cell surface receptor signaling for ‘Normal’ Cluster. Interaction data obtained from String was visualized using Cytoscape.

**Figure S4.**
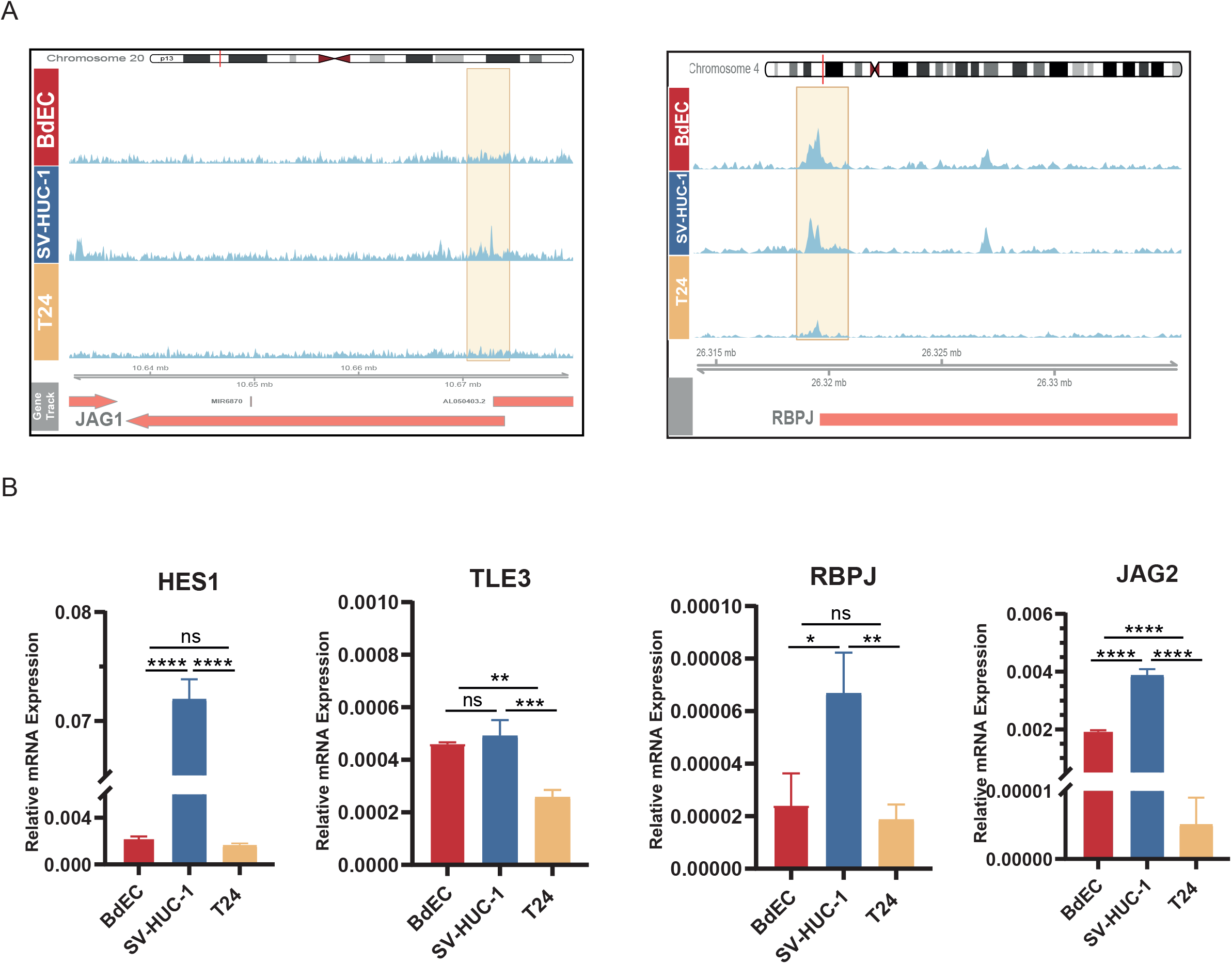
Association of KDM6A with genes involved in Notch signaling. (A) Snapshot images showing KDM6A signal in BdEC, SV-HUC-1, T24 cell lines at JAG1 and RBPJ loci. (B) Barplots showing the result of RT-qPCR analysis performed for *HES1, TLE3, RPBJ, JAG2* genes. Error bars depict the sd of three technical replicates. * pvalue < 0.05, ** pvalue < 0.005, *** pvalue < 0.0005, **** pvalue < 0.0001, ns = not significant

**Figure S5.**
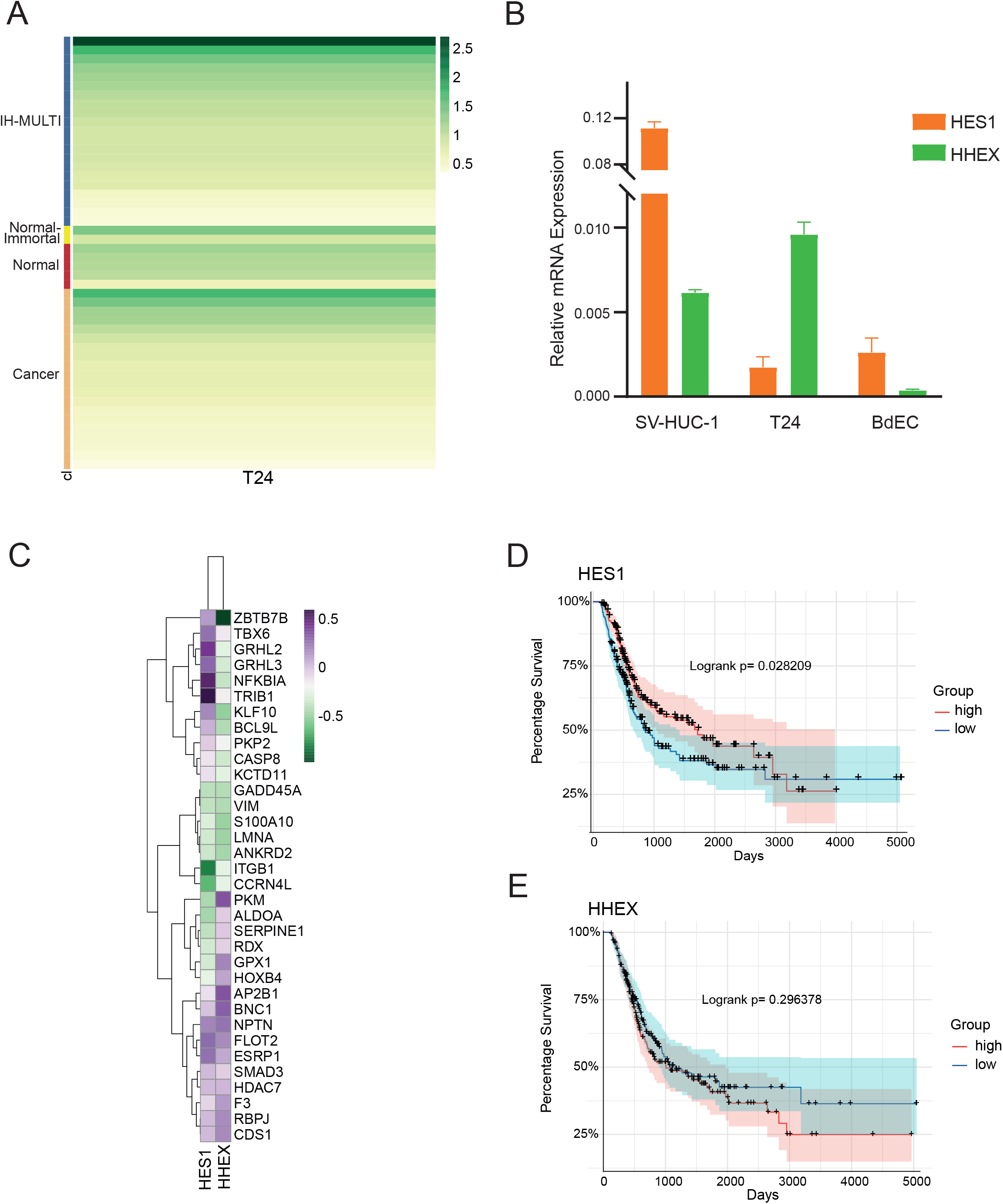
HES1 and HHEX interplay in bladder cancer cell lines and their association with patient survival. (A) Classification of all T24 KDM6A peaks. (B) RT-qPCR expression analysis of *HHEX* and *HES1* in normal and cancer bladder cell lines. Error bars depict the sd of three technical replicates. (C) Heatmap demonstrates the correlation between the expression of *HES1, HHEX* and the genes involved in regulation of developmental process in bladder cancer cell lines. (D-E) Kaplan-Meier graphs show the overall survival of primary bladder cancer patients, grouped according to the expression status of *HES1* (D), *HHEX* (E).

**Figure S6.**
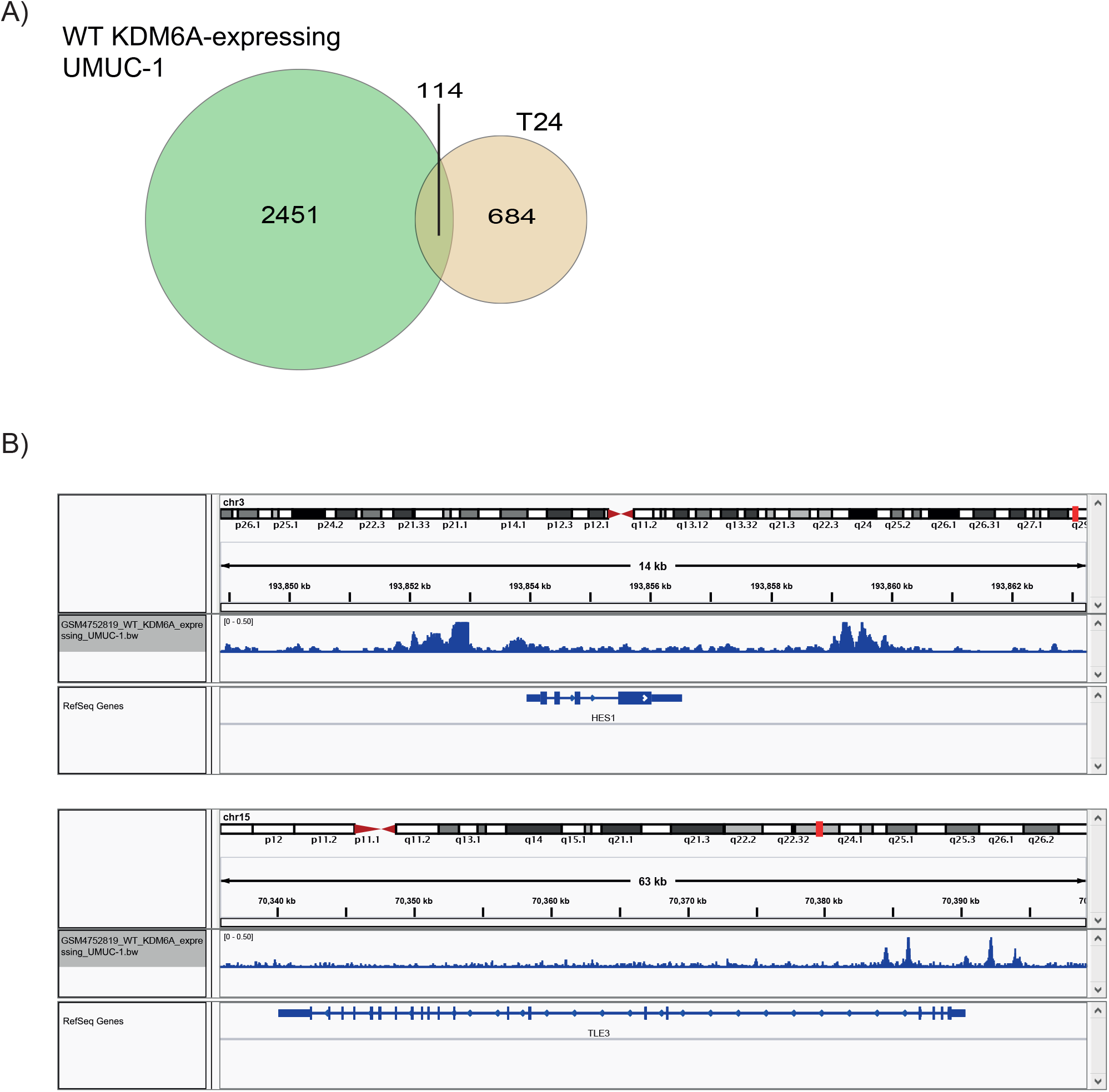
Compar son of KDM6A peaks dent f ed for T24 cell l ne w th the peaks called n WT KDM6A-expressing UMUC-1 cells. (A) Venn diagram shows the KDM6A peak overlap. (B) Snapshot images depict the occupancy of KDM6A at HES1 and TLE3 loci in WT KDM6A-expressing UMUC-1 cells.

**Figure S7.**
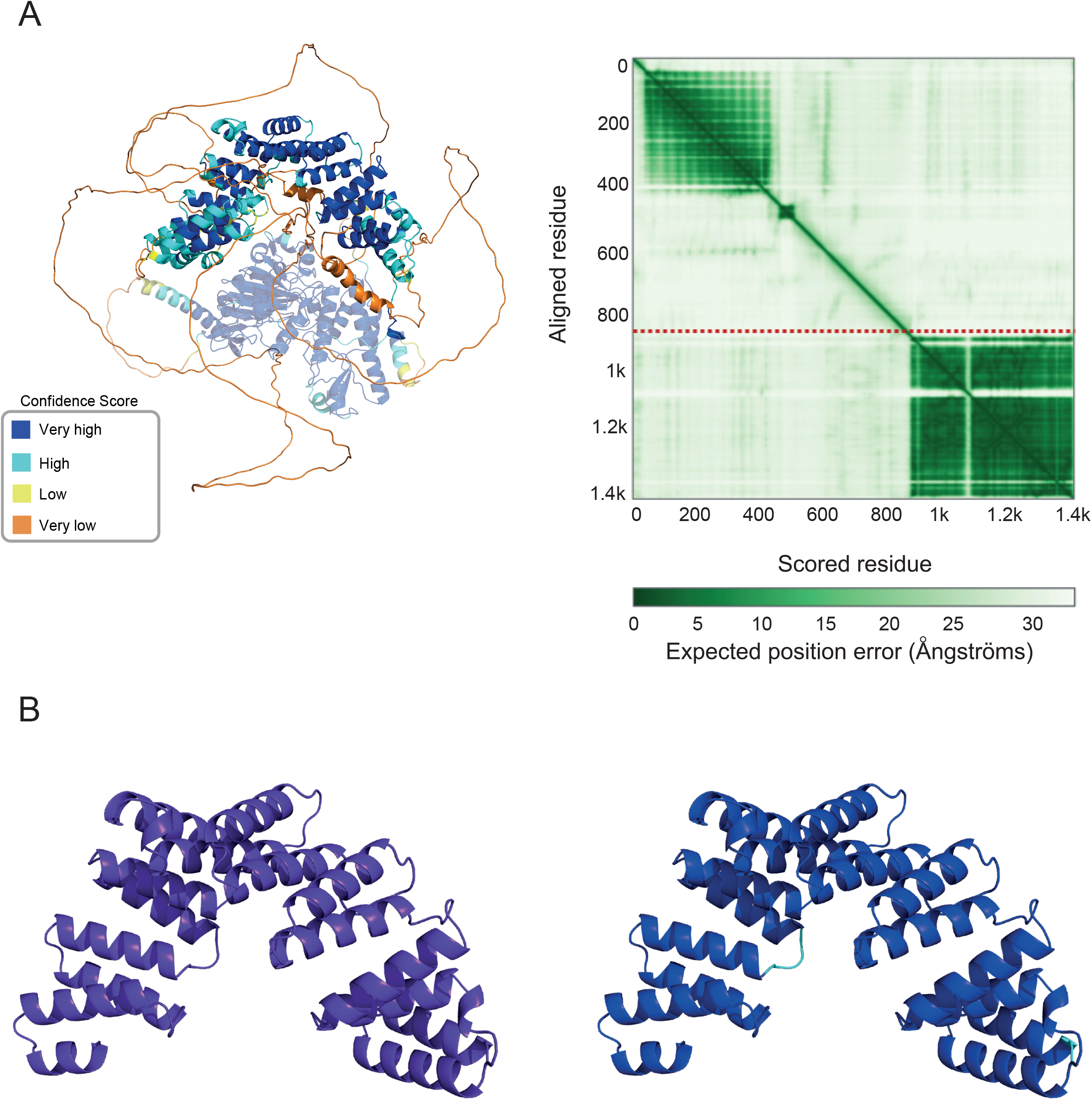
AlphaFold models of KDM6A protein. (A) Full-length KDM6A model taken from AlphaFold Database given in confidence score coloring on the left side (https://alphafold.ebi.ac.uk/entry/O15550). The confidence score decreases gradually towards dark blue, light blue, yellow, and orange. The lost part in the truncated isoform is shown in pale. PAE graph taken from the model’s page is given on the right side together with the red dotted line showing the position of the 895th residue. Estimated position error increases from dark green to white. Darker regions on the diagonal and off-diagonal show well-predicted domain organization and relative domain positions, respectively. (B) AF2 KDM6A-TPR model obtained on a local machine. Confidence score coloring is provided for the best-ranked model (right).

**Figure S8.**
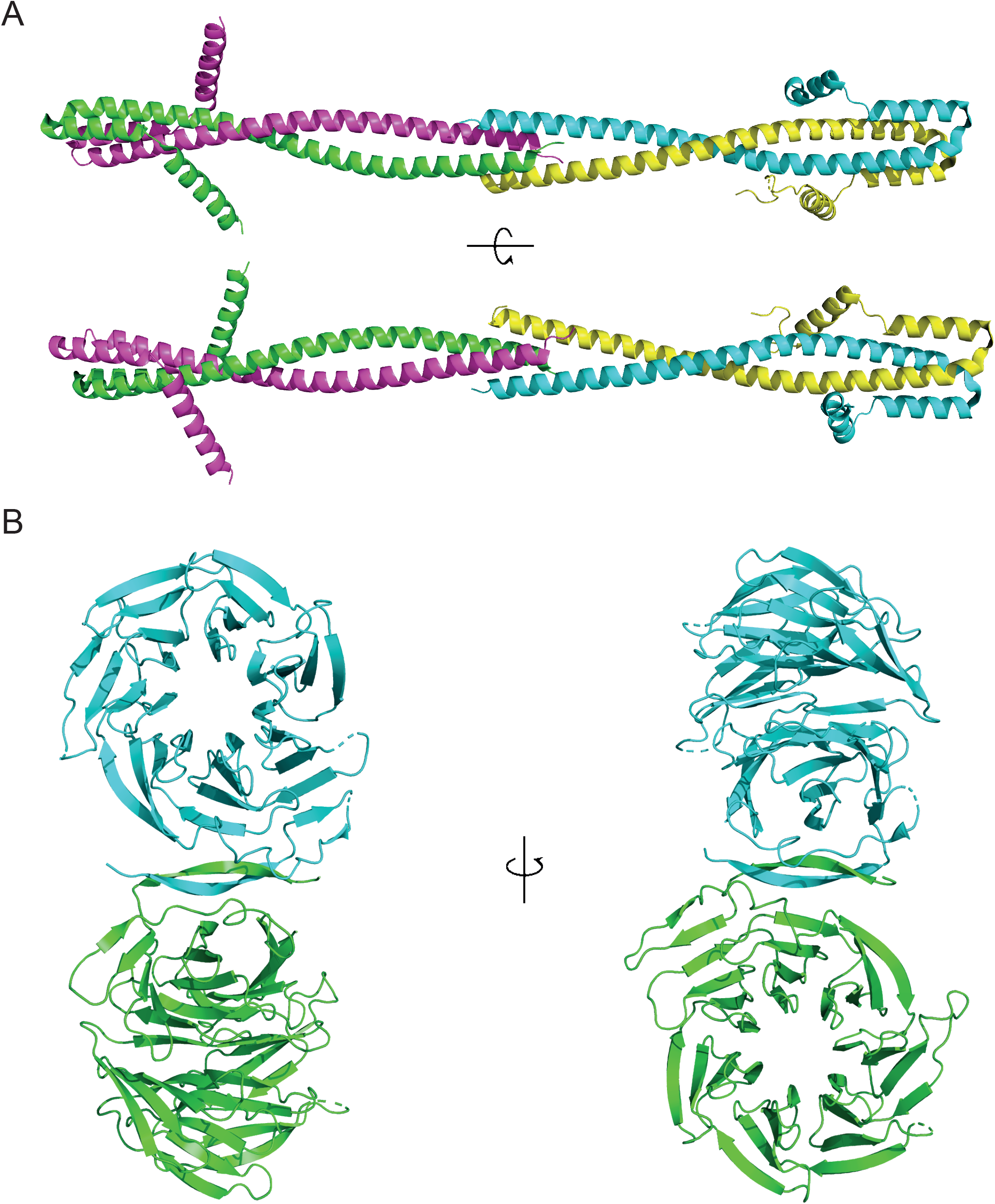
Conserved domain structures in TLE1. (A) N-ter Q domain colored by chains in 4OM3 PDB entry. (B) C-ter WDR domain colored by chains in 1GXR.

**Figure S9.**
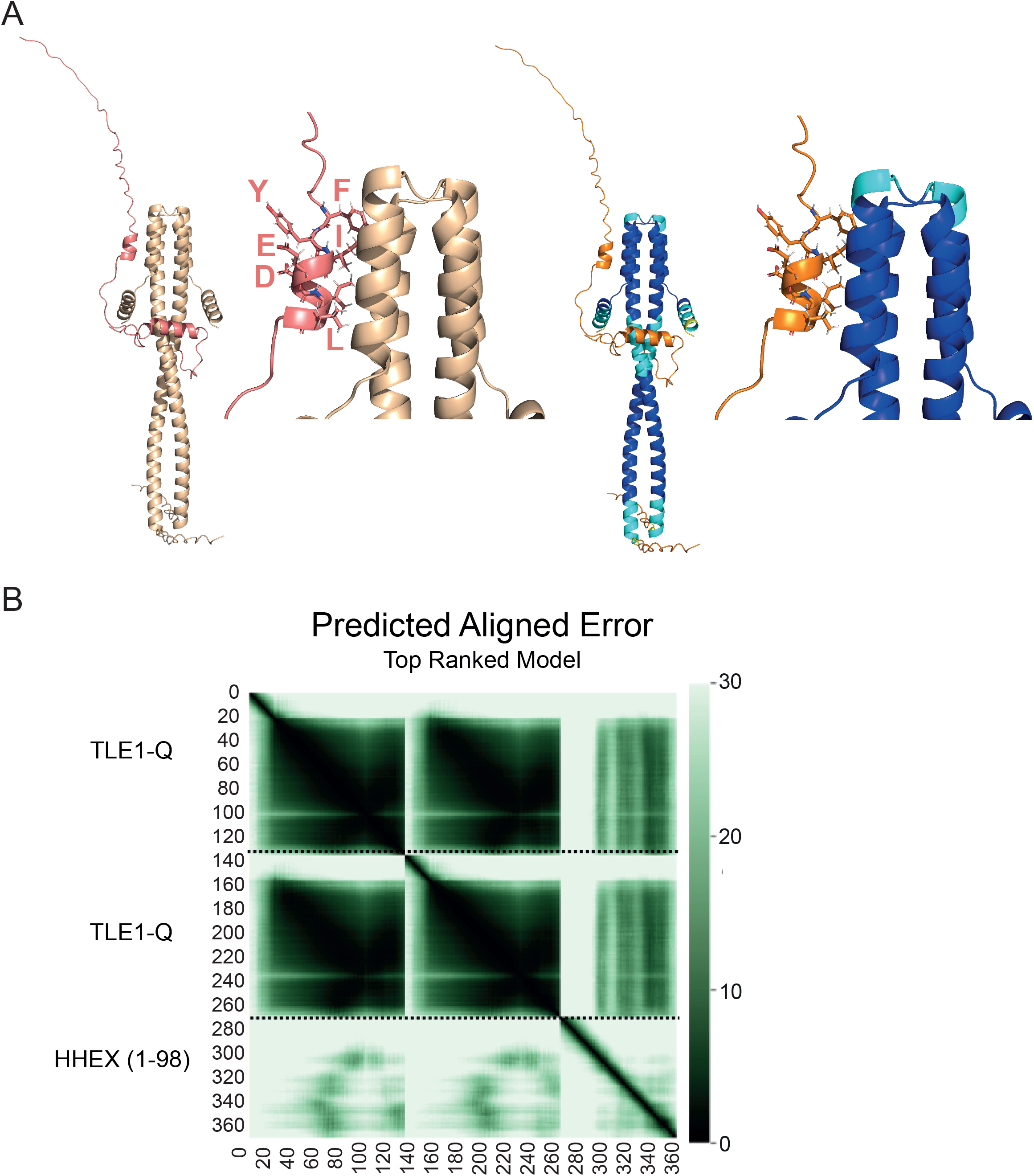
AlphaFold model of TLE1-Q:HHEX complex. (A) The best model obtained by AlphaFold-multimer for TLE1-Q (wheat) and HHEX (salmon). Confidence score coloring is also provided on the left side. (B) Predicted aligned error graph for the best model. The red line represents the monomer boundaries.

